# Application of biophysical methods for improved protein production and characterization: a case study on an HtrA-family bacterial protease

**DOI:** 10.1101/2022.01.27.477556

**Authors:** Michael Ronzetti, Bolormaa Baljinnyam, Ishrat Jalal, Utpal Pal, Anton Simeonov

**Author notes:** **Correspondence:** Anton Simeonov, Utpal Pal.

## Abstract

The high temperature requirement A (HtrA) serine protease family presents an attractive target class for antibacterial therapeutics development. These proteins possess dual protease and chaperone functions and contain numerous binding sites and regulatory loops, displaying diverse oligomerization patterns dependent on substrate type and occupancy. HtrA proteins that are natively purified coelute with contaminating peptides and activating species, shifting oligomerization and protein structure to differently activated populations. Here, a redesigned HtrA production results in cleaner preparations with high yields by overexpressing and purifying target protein from inclusion bodies under denaturing conditions, followed by a high-throughput screen for optimal refolding buffer composition using function-agnostic biophysical techniques that do not rely on target-specific measurements. We use the *Borrelia burgdorferi* HtrA to demonstrate the effectiveness of our function-agnostic approach, while characterization with both new and established biophysical methods shows the retention of proteolytic and chaperone activity of the refolded protein. This systematic workflow and toolset will translate to the production of HtrA-family proteins in higher quantities of pure and monodisperse composition than the current literature standard, with applicability to a broad array of protein purification strategies.

**Statement:** The production of a therapeutically-relevant protein family sensitive to coeluting contaminants is greatly improved by optimized expression and refolding workflow. A miniaturized, high-throughput system supported by a function-agnostic biophysics assay and modified data analysis scripts results in a refolded protein that is highly pure, monodisperse, and retains proteolytic and chaperone activity. This approach has broad applicability towards hard-to-express proteins and proteins sensitive to coeluting species. Additionally, novel methods are presented to characterize protein chaperone activity.

## 1 Introduction

Proteases from the high temperature requirement A family (HtrA) are coded for across the animal kingdom and essential to the survival and infectivity of bacteria and other microbes of concern to public health, making them particularly tractable as next-generation antibiotic targets for drug discovery (Backert, Bernegger et al. 2018, Cho, Choi et al. 2020). HtrA homologs display a wide range of quaternary structures that vary in oligomerization and shape, with a majority of HtrA protein structures yet to be studied (Kim, Grant et al. 2011, Wrase, Scott et al. 2011, Hansen and Hilgenfeld 2013, Thompson, Merdanovic et al. 2014, de Regt, Kim et al. 2015, Schubert, Wrase et al. 2015, Zhang, Wang et al. 2017). All known bacterial HtrA-family proteins form homo-oligomers that shift to higher-order oligomeric structures in response to different signals, most often through allosteric regulation via the PDZ domains (Kolmar, Waller et al. 1996, Pallen and Wren 1997, Clausen, Southan et al. 2002, Jiang, Zhang et al. 2008, Krojer, Sawa et al. 2008, Hansen and Hilgenfeld 2013). When a substrate binds to the regulatory PDZ domain, it causes conformational changes throughout the regulatory loops of the protease, removing blocks from the oxyanion hole and S1 specificity pocket and allowing for proteolytic or holdase-like chaperone activity (Schubert, Wrase et al. 2015). Previous groups have reported on the presence of contaminating peptides that co-purify with and influence the oligomerization of HtrA (Schubert, Wrase et al. 2015, Russell, Tang et al. 2016, Zarzecka, Grinzato et al. 2020). Following up, Bai *et al*. give direct evidence of copurified HtrA substrates by mass spectrometry after a standard native purification of DegQ from *E. coli* culture (Bai, Pan et al. 2011). Purifying HtrA proteins under denaturing conditions would provide an effective means of removing potential HtrA-bound ligands but refolding back into a native structure remains a barrier to effective denaturing purifications.

Purifying functional protein from inclusion bodies presents unique challenges regarding the proper solubilization and refolding of denatured proteins back to their native conformations while still producing acceptable and active yields. While there are numerous commercial kits that are designed to provide a wide range of buffer conditions to screen, it is still up to the end-user to decide how to determine whether a target protein has correctly folded. Conveniently, Biter *et al*. use a differential scanning fluorimetry (DSF) -guided workflow to screen buffers for the refolding process by diluting purified, denatured protein into a matrix of buffer agents and salt concentrations and then monitoring for the presence and magnitude of thermal melting transitions (Amadeo, Andres et al. 2016). This method is especially powerful as the primary readout does not depend on the target protein’s function, providing information on refolding conditions without the requirement for any functional testing. Here, previous DSF-guided refolding (DGR) workflows are greatly improved by the development, miniaturization, and optimization of a 384-well microdilution refolding assay, increasing sample throughput and unfolding data replicates. Additionally, the introduction of a modified analysis of DSF datasets is shown to be better at scoring thermal unfolding curves over the more common single parameters like the aggregation temperature (T_agg_).

The *B. burgdorferi* HtrA, BbHtrA, is a therapeutic target shown to contribute to borrelial dissemination in the host and the pathogenesis of Lyme disease (Kariu, Yang et al. 2013, Kariu, Sharma et al. 2015, Ullmann, Russell et al. 2015, Russell, Tang et al. 2016, Thakur, Bista et al. 2022). Interestingly, *B. burgdorferi* codes for only a single copy of the protein, enhancing its tractability as a drug discovery target (Gherardini 2013, Russell, Delorey et al. 2013, Russell, Tang et al. 2016, Ye, Sharma et al. 2016). Prior works characterizing the BbHtrA protein have shown that the proteolytically-inactive S226A mutant elutes with co-purifying peptides, making for an ideal validation target of the workflow (Russell, Tang et al. 2016). Here, we present a roadmap and protocols for an improved overexpression and denaturing purification of BbHtrA followed by function-agnostic biophysical screening and computational scoring for the optimal refolding conditions (Fig 1). The resulting protein product was then characterized by fluorescent caseinolytic cleavage, thermal shift assay, and light scattering methods, to show that refolded BbHtrA maintains its physiological functions as a protease and chaperone.

**Figure 1.**
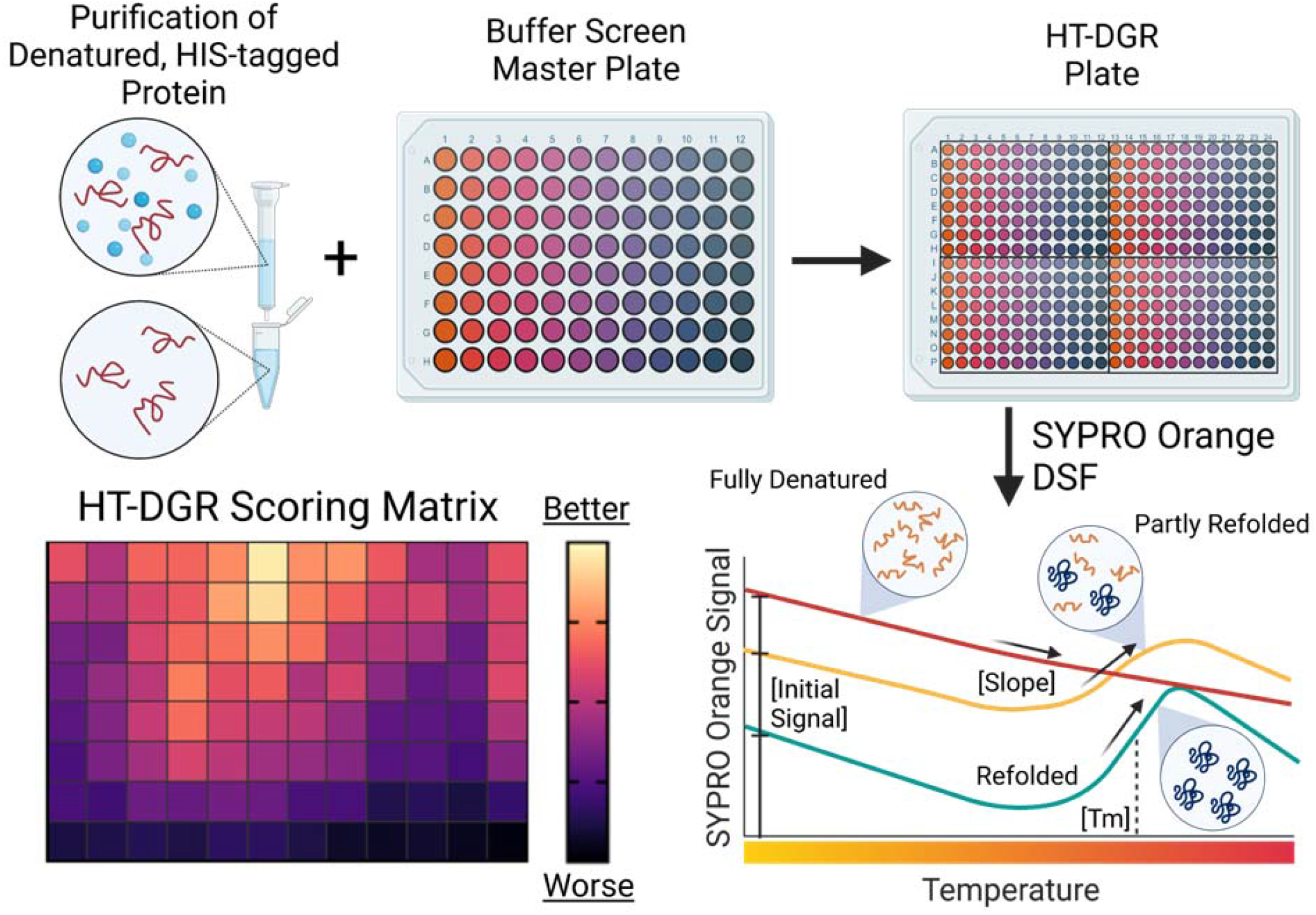
Diagram of the HT-DGR workflow as described in the text. Protein is overexpressed and purified using a denaturing workflow and assayed against a buffer screen of different pH and NaCl concentration, called the HT-DGR plate. Conditions for refolding are tested using differential scanning fluorimetry with SYPRO Orange, providing a biophysical readout of protein unfolding that is used to construct the HT-DGR scoring matrix.

## 2 Results and Discussion

### 2.1 Overview of the method

Starting with overexpressed HIS-tagged protein of interest, a denaturing purification is performed using urea (or an equivalent chaotropic agent) to strip the target protein of co-purifying material, as shown in the method workflow diagram (Fig 1). In parallel, a buffer screen master plate is constructed containing a wide-range of buffer pH, buffer agent, and ionic levels, creating the high-throughput DGR (HT-DGR) assay plate. Denatured protein is dispensed into each well of the HT-DGR plate and refolding takes place through microdilution of the denaturant. After incubation, samples are pulled from each well of the plate and transferred to a 384-well PCR plate to perform a SYPRO-Orange reported differential scanning fluorimetry assay using traditional high-throughput quantitative PCR (qPCR) instrumentation. This DSF data is then parsed and analyzed by an in-house R script to integrate different parameters of the unfolding curve (such as the slope, initial signal, and midpoint of the sigmoidal curve) into a novel DGR score that provides a single parameter used to score the efficiency of refolding conditions. Preparations of the protein then undergo additional target-specific follow-up experiments to analyze the retention of protease and chaperone activity after refolding, specifically with a fluorescence casein proteolysis assay, isothermal dynamic light scattering, and a novel lysozyme chaperone activity assay based on light backscattering.

### 2.2 Cloning and expression of the HtrA protein

The BbHtrA coding sequence (BB_0104 Uniprot #O51131) was first parsed for any signal peptide that may interfere with downstream protein production and purification using the public server, SignalP. Interestingly, the SignalP models present conflicting results, with the HMM model highlighting a signal peptide cleavage site between position 28 and 29, while the neural network mirrors prior literature by detecting a peptide cleavage site between positions 37 and 38 (Russell, Tang et al. 2016). Gene strands coding for the wild type and catalytically-inactive variant (S226A) were synthesized and inserted into pET28A(+)-HIS-TEV plasmid. BL21 (DE3) Star *E. coli* were then transformed according to the manufacturer’s protocol and clones with the highest expression of protein in a traditional IPTG induction were selected to move forward (Fig 2A).

**Figure 2.**
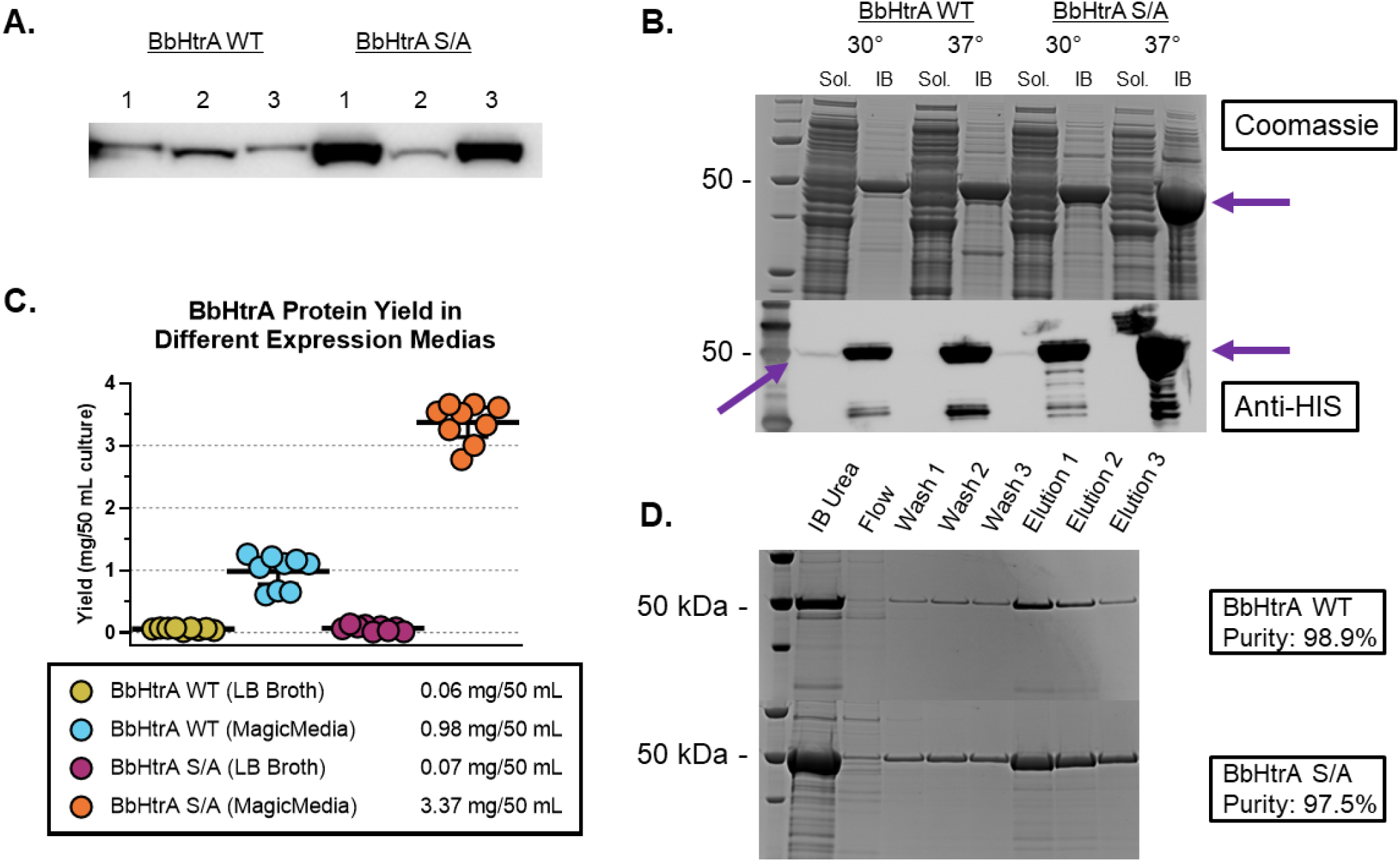
(A) Immunoblot using anti-HIS antibody against three transformation reactions of BbHtrA WT and BbHtrA S/A pET28a constructs. (B) Expression levels of BbHtrA constructs in soluble vs. IB fractions of a 2 mL small-scale culture at different temperatures using the MagicMedia autoinduction system, tested by SDS-PAGE Coomassie staining and immunoblotting. BbHtrA has an expected molecular weight of ~50 kDa, indicated by purple arrows. (C) The protein amount after purification and refolding was measured using the Bradford assay. Expression yields for BbHtrA WT and S/A were improved when using MagicMedia over LB broth, with WT and S/A expressing at 0.98 mg/50 mL (0.78 - 1.18 mg/50 mL 95% CI) and 3.37 mg/50 mL (3.13 - 3.61 mg/50 mL 95% CI) when testing 9 separate 2 mL small-scale cultures at 37 °C for 24 hours. (D) Proof-of-concept workflow of a denaturing purification of BbHtrA proteins results in a highly pure preparation, visualized by SDS-PAGE Coomassie staining.

The biomass and BbHtrA yield was increased by using a high-density autoinduction media, an expression system that is intervention-free after inoculation and improves the yields of target proteins under T7 promoter by using differentially-metabolized carbon sources as compared to traditional multistep LB-IPTG induction (Fox and Blommel 2009, Li, Kessler et al. 2011). Wild-type (WT) BbHtrA and S226A mutant (S/A) were overexpressed in 2 mL MagicMedia autoinduction media at both 30 and 37 °C to optimize protein yield and determine whether the protein is expressed solubly or not. As observed in prior testing with BbHtrA and other HtrA homologs in-house, HtrA proteins tend to express in inclusion bodies (IB) and will shift expression towards the soluble fraction when the temperature is lowered, at the expense of total yield (Fig 2B). The expression of the S/A mutant was also higher relative to the wildtype protein, though the increased target biomass appears almost entirely in the inclusion body fraction (Fig 2B). Testing the target protein yield of 9 individual small-scale cultures (performed in parallel at 37 °C for 24 hours) gave a final yield of BbHtrA WT and S/A at 0.98 mg/50 mL MagicMedia (0.78 - 1.18 mg/50 mL 95% CI) and 3.37 mg/50 mL MagicMedia (3.13 - 3.61 mg/50 mL 95% CI) (Fig 2C), respectively. When compared to a traditional IPTG induction performed in parallel, yields of BbHtrA WT and S/A in MagicMedia were 16.3x and 48.1x higher, respectively (Fig 2C). A pilot denaturing purification of BbHtrA WT and S/A with Co^2+^-NTA spin-columns results in a final purity of 98.9% and 97.5%, respectively, when analyzed by SDS-PAGE and Coomassie staining (Fig 2D).

### 2.3 DSF and dataset analysis for BbHtrA refolding buffer conditions

The HT-DGR plate was constructed by reformatting the 96-well buffer screen master plate into the four quadrants of a 384-well plate using automated pipetting (Apricot Designs PP-384-M) followed by dilution to the final working concentration with ddH_2_O. Next, denatured and purified BbHtrA S/A was dispensed into each well using the Apricot pipetting system to begin the refolding screen. While the method can accommodate a wide-range of conditions and additives to provide project-specific screens, the Hampton Research Solubility and Stability Screen 2 was used as the primary buffer screen because it offers a wide-range pH, buffering agent and NaCl concentration matrix in 96-well format and was used internally with good results in other buffer screening projects. Other commercial kit possibilities include the popular PACT screen, designed to screen for crystallization conditions with common buffers and salts.

After a set incubation period, samples from the BbHtrA S/A HT-DGR plate were tested using a standard SYPRO Orange DSF approach. Briefly, 5 μL of each HT-DGR sample was automatically pipetted (using the Apricot system) into a 384-well PCR plate that was acoustically dry-spotted with 5 nL of SYPRO Orange. After a brief incubation and centrifugation, the PCR plate was run on a Roche LightCycler 480 II and data processed using the Roche Thermal Shift Analysis software. Of note, the BbHtrA S/A refolding plate was incubated overnight at 4 °C after initial readings at 30 and 90 minutes did not produce any DSF signal.

Most thermal shift analysis reports unfolding data as a single parameter signifying the midpoint of the sigmoidal fit of thermal unfolding curves, known as the T_agg_. The results of the BbHtrA S/A HT-DGR screen show reproducible T_agg_ values across the replicate refolding reactions on our plate with only a ~1 °C difference in T_agg_ among the majority of buffer conditions tested (Fig 3A). There was a clear relationship between the pH, salt concentration, and the three parameters calculated from a thermal shift unfolding curve: T_agg_, the initial signal, and the slope of the first derivative at the onset of melting (Fig 3B). Indeed, the BbHtrA protein quaternary structure is known to be sensitive to the concentration of salt in buffer, presenting substrate-independent shifts in oligomeric assemblies at different concentrations of NaCl, which fits well with the apparent sensitivity of the refolding process to salt levels (Russell, Tang et al. 2016).

**Figure 3.**
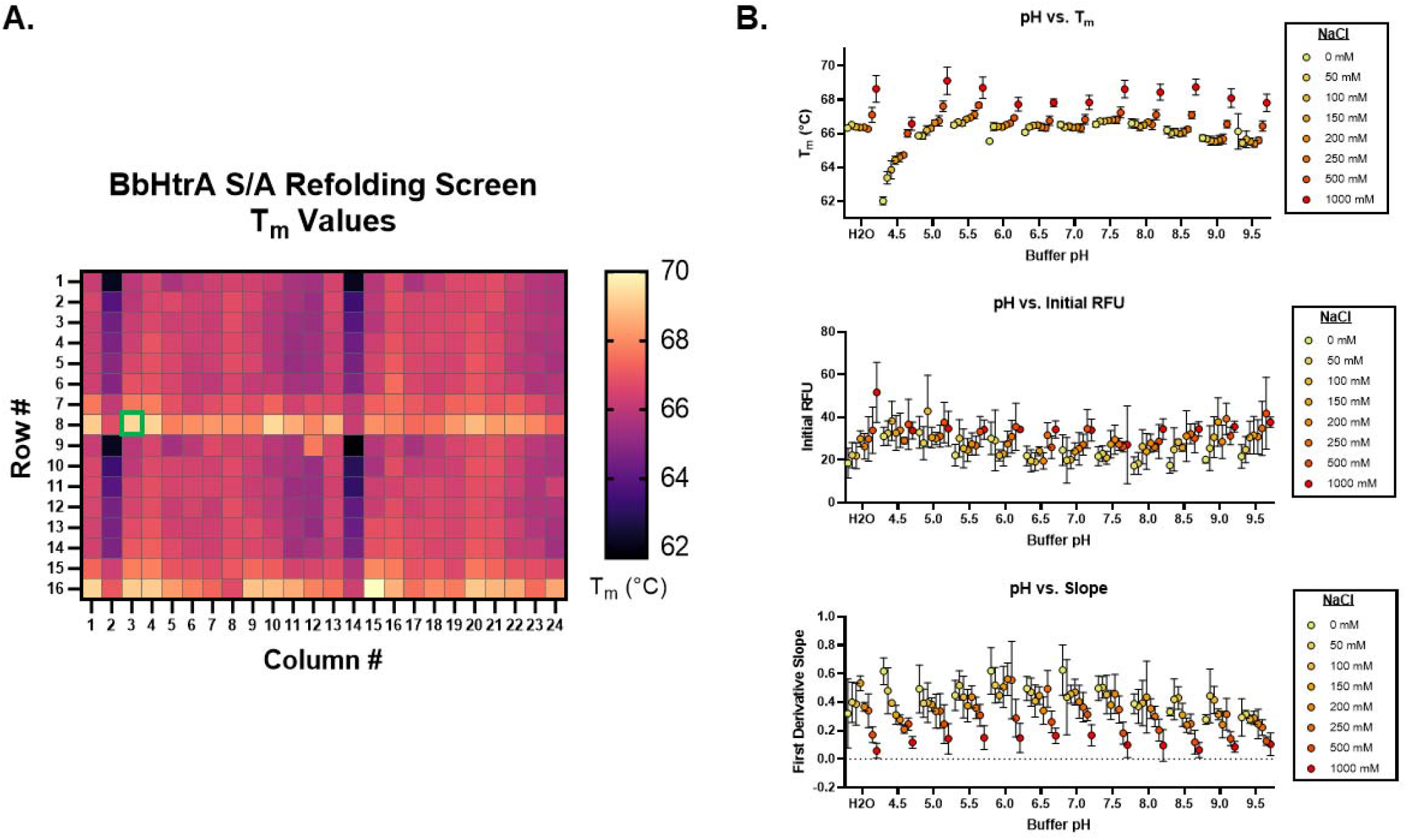
(*A*) Heatmap of T_agg_ values from an HT-DGR screen for the refolding of BbHtrA S/A. The green box is around the sodium citrate tribasic (pH 5.0, NaCl 1000 mM) condition. (C) Values for the T_agg_, initial signal (RFU), and slope of the first derivative plotted against buffer pH, with individual points colored according to the salt concentration. Error bars show the standard deviation for three biological replicates.

Recent work on DGR assays have also analyzed the data by reporting the multiplication of the T_agg_ by the height of the first derivative peak, thereby correlating peak height as a measure of the protein refolding yield (Amadeo, Andres et al. 2016, Yuanze, Niels van et al. 2017). In this work, the ‘DGR score’ is synthesized to describe the refolding stability and yield as represented by the multiplication of the T_agg_ by the maximum slope of the first derivative peak, and then dividing this result by the initial SYPRO Orange signal before the thermal ramp has been applied. The initial signal of a DSF curve when using dyes like SYPRO Orange serves as a convenient indicator of the unfolding status of a sample at baseline. The dye emits fluorescence energy only after binding to hydrophobic regions in unfolded and aggregated protein species, therefore, the initial signal directly correlates with the amount of unfolded, aggregated, or disordered material present in a refolding reaction.

The thermal shift analysis improves after integrating the initial signal and slope of the derivative peak into the DGR score, giving the highest refolding signals with MES, BisTris, Imidazole, and HEPES buffering agent at pH range 6.0 to 7.5 and NaCl concentrations below 100 mM (Fig 4A). The first derivative of the thermal unfolding curves of the top four selected buffers all show single, sharp peaks indicative of a homogenous and folded species, unlike less folded or unfolded populations encountered in the suboptimal buffer conditions (Fig 4B, green curves are optimal conditons, yellow are suboptimal). It is worth noting that an analysis of the HT-DGR screen using T_agg_ as the endpoint would advance sodium citrate tribasic [pH 5.0, NaCl 1000 mM] (Fig 3A, green square) as a top candidate, though the shallow slope and high initial signal in the DSF thermal unfolding curves indicate an unfavorable refolding buffer (Fig 4C).

**Figure 4.**
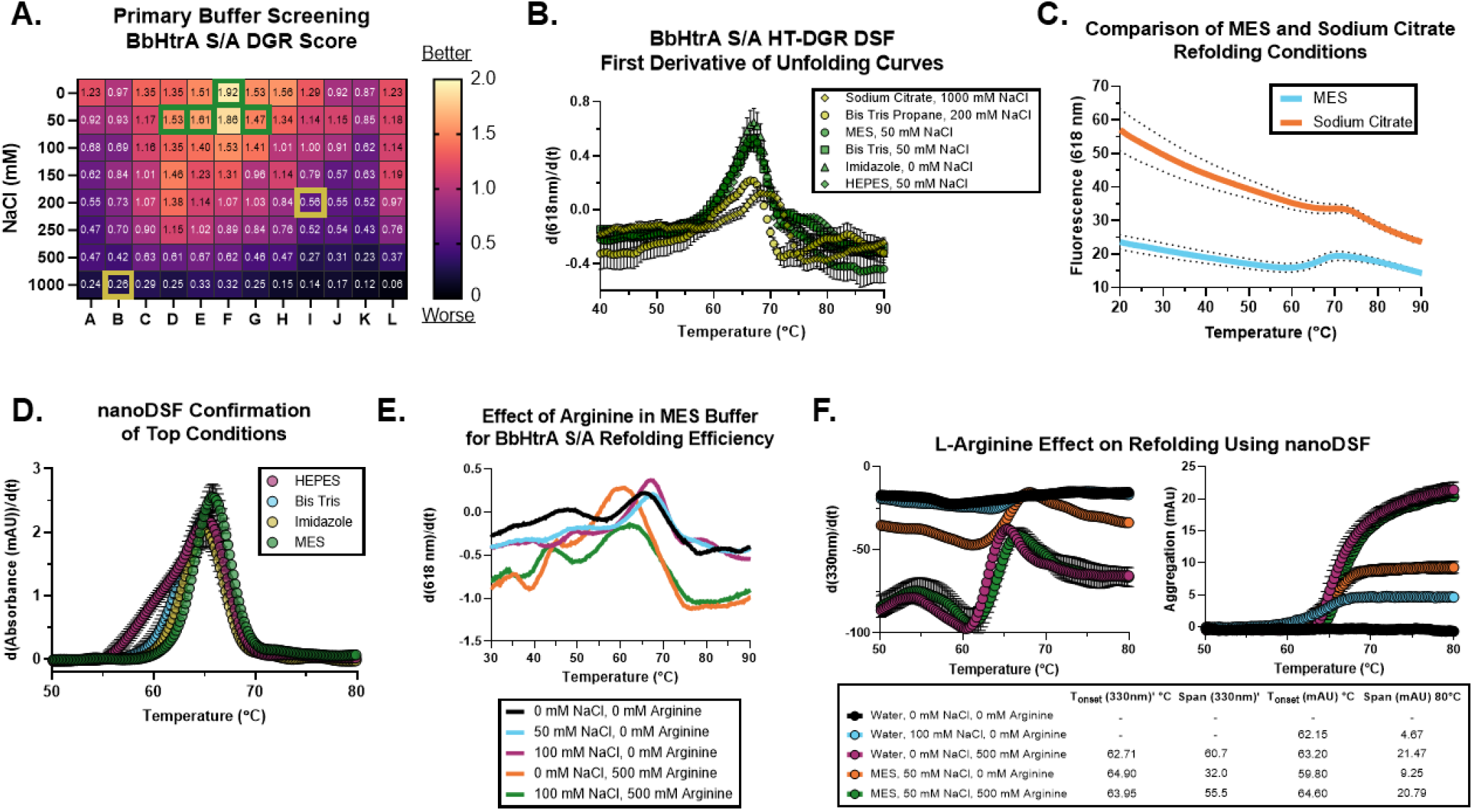
(A) DGR scores for BbHtrA S/A show a more optimal folding reaction at lower NaCl concentrations around 0 - 100 mM and buffer pH between 6.0 – 7.5. The top four conditions are shown with a green box, and representative suboptimal conditions are shown with a yellow box. (B) DSF first derivative unfolding curves for high and low DGR scoring conditions show clear differences in peak height and slope. Note the initial values between different buffering systems indicating different degrees of total unfolded population: Sodium citrate: 55.79 ± 8.19, BisTris Propane: 45.90 ± 7.75, MES: 31.58 ± 6.65, BisTris: 31.58 ± 6.73, Imidazole: 20.87 ± 8.67, HEPES: 33.41 ± 5.75. (C) The raw SYPRO Orange DSF curves for the top condition from the HT-DGR screen (MES) and the top condition when analyzed using Tagg value only (sodium citrate). Data shown is the mean of three replicate biological reactions with dotted lines showing standard deviations. (D) nanoDSF first derivative aggregation curves for the top four conditions of the BbHtrA S/A refolding screen. Data shown is the mean of three replicate biological reactions with error bars showing the standard deviation. (E) The raw SYPRO Orange DSF curves for MES buffer samples with and without arginine. Data shown is the average of three replicate experiments. (F) Comparison of the effects of L-arginine on the refolding efficiency of BbHtrA S/A using nanoDSF. The T_onset_ and span of the 330nm fluorescence first derivative and aggregation readouts are given, showing the improvement in refolding yield in the presence of arginine. Data shown is the average of three replicate experiments with error bars signifying the standard deviation.

As further confirmation of the sample folding status, the thermal stability of BbHtrA S/A in top refolding conditions was analyzed using nanoDSF, an orthogonal DSF method that monitors shifts in both amino-acid autofluorescence and in backscattering aggregation signals during a thermal ramp (Krakowiak, Krajewska et al. 2019, Magnusson, Szekrenyi et al. 2019). While each condition produced strong peaks at ~65 °C, in agreement with T_m_ and T_agg_ measurements, the MES buffer system was chosen to advance as it shows the sharpest, highest peak in the backscattering aggregation curve (Fig 4D).

### 2.4 Probing the effect of L-arginine on refolding reactions

The amino acid L-arginine is often included as an additive in refolding buffers to deter aggregation of target proteins by reducing disordered protein-protein interactions along the refolding process (François, François et al. 2004, Baynes, Wang et al. 2005, Yuanze, Niels van et al. 2017). In order to potentially support higher refolding yields, L-arginine was introduced after primary buffer screening at high-millimolar working concentrations. Although DSF signals have been established for other protein targets using L-arginine in the literature, it appears to interfere with the SYPRO Orange thermal shift readout in our system, so the alternative nanoDSF technique was applied to samples including L-arginine as described before (Fig 4E). As expected, there was no signal in the negative control water-only condition, indicating an absence of properly folded protein in the sample. The inclusion of 500 mM arginine in water produced a significant signal, while only minimal signal was apparent when only NaCl was present (Fig 4F). The MES buffer system was markedly improved with the addition of 500 mM arginine, which induced a significant shift in signal span for both intrinsic amino acid fluorescence at 330 nm (from 32.0 to 55.5) and the backscattering sensor for sample aggregation (from 9.25 to 20.79) (Fig 4F). Notably, while the increase in signal span indicates more properly folded protein contributing to unfolding signals, the presence of arginine did not significantly shift the T_agg_ or T_onset_, suggesting that arginine enhances refolding through a mechanism that does not affect thermal stability of the protein and in agreement with previous work on the role of arginine in protein refolding (Baynes, Wang et al. 2005).

As a final test of the optimized MES buffer system with arginine, the refolding reaction was scaled up to 1 mg of purified BbHtrA WT and S/A diluted to the same concentration as the buffer screen (0.25 mg/mL) and then dialyzed overnight into the final refolding buffer (50 mM MES, pH 6.5, 50 mM NaCl, 500 mM L-arginine). In parallel, native BbHtrA WT and S/A was produced as a control using cell-free protein expression systems following the manufacturer’s suggested protocol with 500 ng plasmid DNA in 50 μL reaction volume. In-house experiments and prior literature on BbHtrA have shown PBS (pH 7.4) to be the primary buffer for activity and storage, further supported by isothermal DLS data indicating that BbHtrA S/A is monodisperse in PBS but exhibits polydispersity in MES refolding buffer (Fig S1A) (Coleman, Toledo et al. 2016, Zhang, Qin et al. 2019). Following buffer exchange of the two preparations to PBS, the protein was analyzed using nanoDSF as described earlier. There was no significant difference in the T_agg_ of BbHtrA produced by cell-free conditions or the scaled-up refolded reaction, interpreted as confirmation of an optimal refolding of the BbHtrA proteins as a result of the HT-DGR-guided refolding buffer design (Fig 5A).

**Figure 5.**
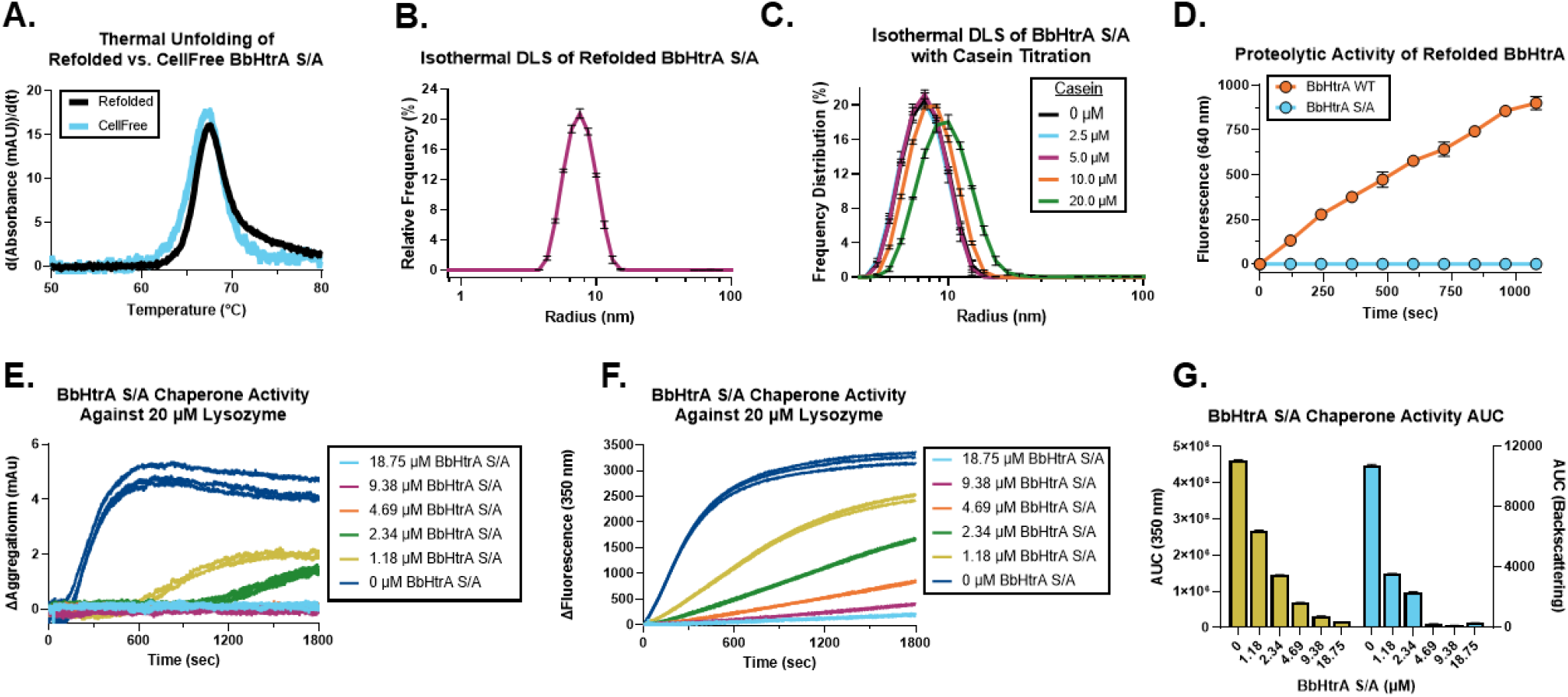
(A) Comparison of the thermal unfolding profiles of BbHtrA S/A expressed using cellfree native conditions and optimized refolding conditions show similar thermal unfolding profiles. (B) Isothermal DLS of the final refolded BbHtrA S/A product in PBS. Data shown is the mean % relative frequency distribution by *r_H_* across three replicates, with error bars showing the standard deviation. (C) Isothermal DLS of BbHtrA S/A in the presence of a casein titration. Data shown is the mean % relative frequency distribution by *r_H_* across three replicates, with error bars showing the standard deviation. (D) The protease activity of BbHtrA WT and S/A after refolding was tested using a casein-BODIPY substrate. Data shown are three independent biological replicates, with error bars showing standard deviations. (E-G) Detection of chaperone activity of refolded BbHtrA S/A with lysozyme using nanoDSF. Graphs show the change in (E) aggregation and (F) 350 nm fluorescence of TCEP-induced lysozyme unfolding in the presence of a titration of BbHtrA S/A. (G) Area under the curve (AUC) values for the aggregation and 350 nm fluorescence readings from the chaperone activity test. The figure shows the mean AUC value for three biological replicates with error bars showing the standard deviation.

The homogeneity of the refolded BbHtrA S/A product was further characterized using isothermal dynamic light scattering to check the polydispersity index (PDI) and hydrodynamic radius (*r_H_*) of the sample. Using the DLS functionality of the Prometheus Panta system (NanoTemper Technologies) to perform an isothermal hold at 37 °C, refolded BbHtrA S/A protein in PBS shows a cumulant *r_H_* of 7.47 nm ± 0.02 with a PDI of 0.06 ± 0.01, indicating a highly monodisperse population of protein (Fig 5B). We also sought to test the ability of BbHtrA S/A to oligomerize in response to substrates in solution. When BbHtrA S/A was incubated with a titration series of casein, the cumulant *r_H_* increases with higher casein concentrations (PBS: 7.47 nm ± 0.02, 20 μM casein: 9.21 nm ± 0.04), demonstrating the formation of higher order oligomeric structures (Fig 5C).

### 2.5 Characterizing refolded protein activity

Further confirmation of proper BbHtrA refolding was observed by analyzing the proteolytic cleavage of a casein-BODIPY substrate by wildtype BbHtrA, produced using the above established BbHtrA refolding protocol. Briefly, 50 nM of refolded BbHtrA WT or S/A was mixed with 10 μg/mL casein-BODIPY in a 50 μL reaction and immediately read in kinetic mode for 20 minutes, where the increase in fluorescence when incubated with BbHtrA WT, but not catalytically-inactive BbHtrA S/A, indicates the proteolytic liberation of fluorescent peptide fragments from the casein-BODIPY substrate (Fig 5D).

The ability of the HtrA protein to prevent lysozyme denaturation in the presence of denaturant is a well-established method of analyzing chaperone activity (Zarzecka, Modrak-Wojcik et al. 2018, Zarzecka, Harrer et al. 2019, Zarzecka, Matkowska et al. 2021). The current microplate-based absorbance measurements used for this experiment require large sample volumes (up to 1 mL) and high concentration of protein (up to 50 μM) in order to detect aggregation events. We hypothesized that nanoDSF instrumentation would present a more sensitive detection scheme for monitoring lysozyme unfolding and produces additional data from the intrinsic fluorescence detection channels. Indeed, the 350 nm fluorescent channel was more sensitive to lysozyme unfolding at lower concentrations of TCEP as compared to the backscattering aggregation channel (Fig S1B/C). The addition of refolded BbHtrA S/A showed a concentration-dependent reduction in the unfolding signals of lysozyme samples in both backscattering and 350 nm fluorescence, indicative of the retention of holdase-like chaperone activity (Fig 5E-G). Again, the 350 nm fluorescence channel was a more sensitive measurement of the reduction in lysozyme unfolding than backscattering absorbance readings, as the reduction in unfolding signal remains linear throughout the BbHtrA S/A titration series (Fig S1D). As a control for the nanoDSF bulk measurement of all proteins in solution, the contribution of additional protein to the sample does not seem to interfere with the intrinsic fluorescence shifts observed by lysozyme unfolding, as unfolding signals are still apparent when lysozyme is in the presence of negative control protein luciferase, which has no reported chaperone function, but ablated in the presence of refolded BbHtrA S/A (Fig S1E/F). Importantly, there was no effect of TCEP on the unfolding of BbHtrA S/A or change in the BbHtrA signal at the temperature used for chaperone activity experiments (40 °C) as demonstrated in a temperature-stepping experiment (Fig S1G).

### 2.6 Automating the Analysis of Refolding DGR Scores

The high-throughput format of the HT-DGR assay and its increased information content required the use of scripted analysis to guide decision-making on protein refolding scores. Additionally, there was no currently available DSF analysis software able to derive the DGR score as presented above. To that end, an automated script was written in R to proceed directly from data files describing the thermal unfolding curves all the way through DGR analysis. Using data directly from the Roche TSA program (or any similar thermal shift curve analysis), the script pulls and calculates the T_m_, initial signal, slope, and DGR parameters for every sample. User-supplied plate maps then link sample wells to individual buffer conditions and enable graphical output of the thermal curve parameters and DGR score for each buffer component, as well as information related to refolding score trends in the data (Fig S2). The script is available on GitHub (https://github.com/mronzetti/HT-DGR) along with use instructions.

## 3 Conclusions

The Anfinsen hypothesis states that all necessary information for protein folding is contained in the primary peptide sequence, and so it follows that if one can find the most stable version of the protein that it must represent the native folding state. On this hypothesis, this study presents a roadmap to significantly improving the yield and purity of HtrA (and related) proteins using a widely-accessible high-throughput assay to rapidly screen buffer conditions for optimal refolding of a target protein, all without dependence on any functional assay. While most methods of refolding protein involve long dialysis periods or the use of a multitude of spin columns, the microdilution method presented above allows for rapid screening of conditions in a fraction of the time, generating readily interpretable data on protein refolding. In addition to establishing a miniaturized, systematic, and efficient protein refolding procedure, we address and improve upon the lack of reliable biophysical methods to rapidly detect the formation of correctly refolded proteins, especially those related to the HtrA protein family.

Spurious thermal melts are relatively common in DSF experiments, but by miniaturizing the assay into 384-well format and automating the liquid handling processes, we increase the number of replicates as compared to previous methods and reduce the error in protein and dye concentrations that is inherent to human pipetting. Large errors in pipetting, encountered more as the plate density increases, can alter the entire outcome of the primary refolding buffer screening and obfuscate optimal conditions. Additionally, by testing conditions from the primary buffer screen with a complementary thermal shift approach, more data is revealed on favorable refolding conditions that would otherwise not be detectable with the SYPRO Orange readout. The flexibility of the HT-DGR format allows researchers to finetune the buffer conditions and concentrations tested with target-dependent changes without interfering with assay design.

The reduction of error also unlocks analysis of the initial signal parameter of the thermal shift melt, often discarded or masked by normalizing melt curves before analysis. This allows us to introduce the DGR scoring parameter that is better able to describe the refolding efficacy of a sample over single parameter statistics commonly used like T_m_. Evidence of the improvement in scoring is clear when comparing refolding buffer candidates for BbHtrA that would advance from either T_agg_ or DGR endpoints. Further, the creation of a script to handle all thermal unfolding analysis automatically greatly reduces researcher handling time and improves consistency of the end result.

The catalytically-inactive form of HtrA proteins, often used for structural and biophysical characterization work, are unable to digest copurifying peptides; as such, native preparations of these targets will shift the population away from an apo-form towards a mixture of activated states. Through the use of new dynamic light scattering technology, the population of refolded BbHtrA S/A was shown to be highly monodisperse, suggesting that the denaturing purification was effective in removing contaminating species from our HtrA protein. Previous groups isolating BbHtrA S/A revealed a mixture of monomers, trimers, hexamers, and higher order structures after purification. To the best of our knowledge, the present refolding workflow has produced the first and only example of a highly pure, active, and monodisperse BbHtrA S/A product. Importantly, this homogenous preparation of HtrA protein also displayed an ability to oligomerize in response to an appropriate substrate, further evidence of proper BbHtrA refolding and validation of isothermal DLS as a powerful and material-efficient manner of monitoring HtrA oligoermization shifts in solution.

Characterization of HtrA holdase-like chaperone activity is often accomplished by monitoring changes in the rate of denatured lysozyme aggregation in the presence or absence of HtrA. Aggregation of lysozyme in the presence of a disulfide-bond denaturant results in turbidity over time that can be detected by western blot or light scattering measurements, but these assays as described in the literature are low sensitivity and material-inefficient. There is ample literature supporting the use of nanoDSF as an orthogonal thermal shift method for monitoring protein stability, but this workflow represents the first use case for monitoring chaperone activity with the lysozyme model substrate. In addition, the use of tryptophan intrinsic fluorescence shift was shown to be more sensitive to the unfolding of lysozyme as compared to traditional light scattering methods, minimizing the necessary concentration of chemical denaturant needed and potentially enabling chaperone activity assays with denaturant-sensitive chaperone proteins.

## 4 Materials and Methods

### 4.1 Cloning and Expression

The BbHtrA amino acid sequence (Uniprot O51131) was parsed using the SignalP 3.0 signal peptide detection program with both the hidden Markov model (HMM) and neural network modes, and the resulting signal peptide detected was omitted from the final sequence. The BbHtrA gene strand (AAC66500, 1326 bp) was then cloned into the pET28a (+)-TEV vector using the NdeI/BamHI restriction sites, retaining the N-terminal 6XHIS tag and TEV protease site for downstream tag cleavage. Constructs were verified at the T7 promoter for correct insertion, after which BbHtrA WT and the inactive S226A point mutant (S/A) were transformed into BL21 Star (DE3) chemically-competent *E. coli* which contain a rne131 mutation that leads to higher mRNA stability and thus higher-level protein expression over standard BL21 strains.

For the transformation reaction, 10 ng of plasmid in 5 μL ddH_2_O was mixed gently with one vial of BL21 Star cells and transformed following the manufacturer’s standard protocol (One Shot BL21 Star (DE3), ThermoFisher). 80 μL of the reaction was streaked onto kanamycin agar plates and incubated at 37 °C overnight, after which the protein expression of three separate colonies was checked with a small-scale culture according to the manufacturer’s protocol (2 mL culture of Terrific Broth at 37 °C overnight, 50 μg/mL kanamycin, 0.5 mM IPTG). The culture was spun down (2,000 × g, 10 min, 4 °C) and lysed using 250 μL BugBuster Master Mix for 30 min at room temperature (RT). The crude lysates were spun down at 20,000 × g for 30 min at 4 °C, after which 5 μL of each lysate (1:50^th^ of the original culture) was separated using SDS-PAGE and analyzed by immunoblotting for the histidine tag (1:2500 α-HIS). For expression using the MagicMedia expression medium (ThermoFisher), a starter culture at 1/20^th^ the final culture vessel volume was prepared using stabs of single colonies form freshly streaked plates in LB media with 50 μg/mL kanamycin and incubated at 37 °C overnight with vigorous shaking (250 rpm). The whole starter culture was then added to pre-warmed MagicMedia with 50 μg/mL kanamycin, and then incubated at 37 °C overnight with vigorous shaking (250 rpm).

### 4.2 Protein Purification

E. coli expressing BbHtrA constructs were spun down at 2,000 × g at 4°C for 10 minutes, after which the cell pellets were resuspended in BugBuster Master Mix at 8 mL BugBuster/ 50 mL culture volume and then mixed with a tube inverter for 30 minutes at room temperature. Insoluble cellular debris and inclusion bodies were separated by centrifuging at 16,000 x g at 4°C for 20 minutes, after which inclusion bodies were isolated from the pellet according to manufacturer’s protocol (BugBuster Master Mix, Novagen). Purified IBs containing BbHtrA were resuspended in Denaturing/Binding buffer (20 mM phosphate buffer, pH 7.5, 5 mM KCl, 500 mM NaCl, 8M urea, 20 mM imidazole) at a concentration of 100 mg IB/mL, shaken at RT for 1 hour, and centrifuged at 20,000 x g for 10 minutes at RT. The denatured protein was then purified in 0.2 or 1.0 mL Co-NTA spin columns using a denaturing purification according to manufacturer’s protocol (HisPur Spin Column, ThermoScientific). The concentration of purified protein was determined using a Bradford protein assay with BSA as a standard.

### 4.3 High-Throughput DSF-Guided Refolding

The Hampton Research Solubility and Stability Screen 2 (Hampton Biosciences) was used at the recommended dilution of 4X when constructing the HT-DGR screen, resulting in final concentrations of buffering agent and NaCl of 50 mM and 0-1000 mM, respectively. The 96-well master plate was compressed into the four quadrants of a 384-well nonbinding plate (Cat #) using an Apricot PP-384 pipettor by dispensing 10 μL of each well from the master plate into 28 μL of ddH_2_O, making a 4X dilution in a final volume of 40 μL after subsequent protein dispensing. Purified BbHtrA S/A was first diluted to 5 mg/mL in denaturing buffer before 2 μL of protein was dispensed into each well of the 384-well plate for a final protein concentration of 0.25 mg/mL (5 μM). The plates were then sealed with DMSO-resistant foil and placed on a shaker at 100 rpm at 4 °C. Samples taken at 30 and 90-minute incubation periods gave low and noisy signal, which prompted an extension to an 18-hour incubation to allow for a more complete refolding.

### 4.4 Thermal Shift Analysis

#### 4.4.1 Differential Scanning Fluorimetry

The DSF assay plate was constructed by dry-spotting 5 nL of SYPRO Orange with an acoustic dispenser (Echo 555, Labcyte) into a 384-well PCR plate. The plate was spun down at 200 x g for 1 minute before dispensing 5 μL of the refolding reaction per well and mixing using an Apricot PP-384 pipettor. The plate was sealed, spun down, and immediately run on a Roche LightCycler 480 II using standard DSF conditions (20-95 °C, 0.13 °C/second, 4 acquisitions per °C). The resulting thermal shift data set was then analyzed using Roche Thermal Shift Analysis software to derive target unfolding parameters, namely the first derivative curves, T_agg_ values, slope of the derivative peak, and initial signal before heating, using standard program parameters. The data from the Roche Therma Shift Analysis software and HT-DGR score was parsed and calculated using an in-house script in R (GitHub repository available at: https://github.com/mronzetti/HT-DGR).

#### 4.4.2 Nano Differential Scanning Fluorimetry

Following standard nanoDSF experimental design, 5 μL of refolded protein was loaded into standard capillaries and onto the capillary tray of a Prometheus NT.48 instrument (NanoTemper Technologies) and run using a 5.0 °C/min thermal ramp at 50% LED excitation. The data for each channel was analyzed using the NanoTemper NT.TimeControl software to derive T_agg_ and T_onset_ for each condition.

### 4.5 Protein Activity Characterization

#### 4.5.1 Isothermal Dynamic Light Scattering

Isothermal DLS scans (10 acquisitions, 5 s each, 57% LED intensity, 100% DLS Laser intensity) were performed at 37°C for refolded BbHtrA S/A (10 μM, PBS) in the presence or absence of a casein titration series (20, 10, 5, & 2.5 μM). Samples were incubated for 10 mins, followed by centrifugation (10 min, 4°C, 13000 rpm), filled into Prometheus high-sensitivity capillaries (10 μL each, n=3) and transferred to a Prometheus Panta instrument (NanoTemper Technologies). DLS analysis of BbHtrA S/A *r_h_* and PDI were determined using Prometheus Panta’s size analysis function.

#### 4.5.2 Casein-BODIPY Proteolysis Activity

Protease activity was profiled using a casein substrate that has been labeled in molar excess with BODIPY TR-X dye (EnzChek Protease Activity Kit). First, 16 μL of 62.5 nM BbHtrA WT (50 nM BbHtrA WT final concentration) was added into Greiner 384-well black plates (782096) and spun down. Then, 4 μL of 25 μg/mL casein-BODIPY (5 μg/mL casein-BODIPY final) was added to each well, mixed, spun down, and immediately read on a Tecan Infinite M1000 in kinetic mode for 20 minutes with the following instrument settings: excitation wavelength = 590 nm ± 10; emission wavelength = 645 ± 20; gain = 95. The increase in fluorescence when incubated with active protease indicates the proteolytic liberation of fluorescent peptide fragments from the casein-BODIPY substrate, and proteolytic digestion rates were normalized to their zero timepoint.

#### 4.5.3 Lysozyme Chaperone Activity using nanoDSF/DLS

The chaperone activity of the refolded HtrA protein was tested by monitoring the unfolding of lysozyme in the presence of a chemical denaturant, TCEP, using the nanoDSF Prometheus NT.48 instrument (NanoTemper Technologies). For the isothermal hold experiments, the machine ramps up from a starting temperature of 25 °C to 40 °C at maximum speed and held for 30 minutes, with excitation LED energy set to 50%. For temperature stepping experiments, the machine ramps up from a starting temperature of 25 °C to 37 °C at maximum speed, then ramps up in Δ3 °C steps with a three-minute hold at each point.

For isothermal experiments looking at the concentration-response effect of TCEP on lysozyme unfolding, a 1000 μM 1:2 4-pt titration of TCEP was constructed in PBS, mixed with 20 μM lysozyme, and immediately loaded into standard capillaries and onto the capillary tray of the Prometheus NT.48 instrument. In isothermal experiments looking at the unfolding of lysozyme in the presence of BbHtrA or luciferase, the TCEP concentration was a constant 250 μM, added to a mixture of 20 μM lysozyme and 10 μM of either refolded BbHtrA S/A or luciferase, and read as described above. Experiments analyzing the concentration-response of BbHtrA S/A involved an 18.75 μM 1:2 5-pt titration series of BbHtrA S/A constructed in PBS, mixed with 20 μM lysozyme and 250 μM TCEP, and read in an isothermal hold experiment as described above.

## Abbreviations and Symbols

DSF: Differential scanning fluorimetry
DLS: Dynamic light scattering
HtrA: High temperature requirement A protein
HT-DGR: High-throughput DSF-guided refolding
PDI: Polydispersity index
*r_H_*: Hydrodynamic radius

## 5 Author Contributions

<u>MR</u>: Conceptualization, Methodology, Experimentation, Validation, Analysis, Writing, Review & Editing <u>BB</u>: Conceptualization, Methodology, Validation, Writing, Review & Editing <u>IJ</u>: Methodology, Experimentation, Review & Editing <u>UP</u>: Supervision, Review & Editing <u>AS</u>: Conceptualization, Methodology, Supervision, Review & Editing

## 8 Figure Legends

**Supplementary Figure 1.**
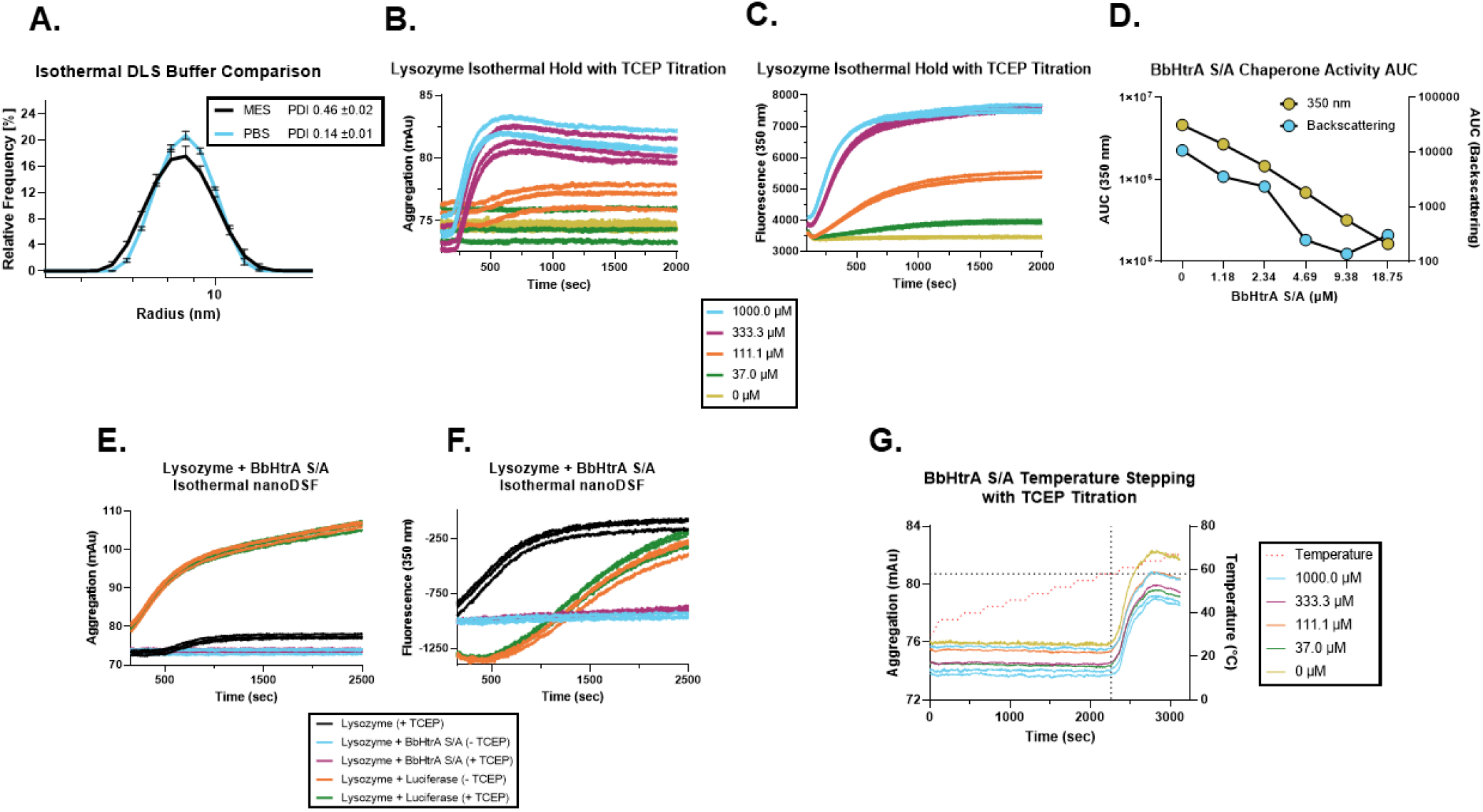
(A) Isothermal DLS of BbHtrA S/A in refolding (MES) or activity/storage buffer (PBS). Data shown is the mean % relative frequency distribution by *r_H_* across three replicates, with error bars showing the standard deviation. (B-C) Raw curves from the (B) backscattering and (C) 350 nm fluorescence channels of a 40 °C isothermal hold of lysozyme in the presence of a titration of TCEP. Three biological replicate traces are shown. (D) Comparison of AUC values from the backscattering and 350 nm fluorescence channels for the BbHtrA S/A concentration-response chaperone activity experiment. Individual points represent the mean of three replicates. (E-F) Raw curves from the (E) backscattering and (F) 350 nm fluorescence channels of a 40 °C isothermal hold of lysozyme and target proteins in the presence or absence of TCEP. Three biological replicate traces are shown. (G) Raw backscattering curves from the Δ3 °C temperature stepping of BbHtrA S/A in the presence of a titration of TCEP. Three biological replicate traces are shown.

**Figure S2.**
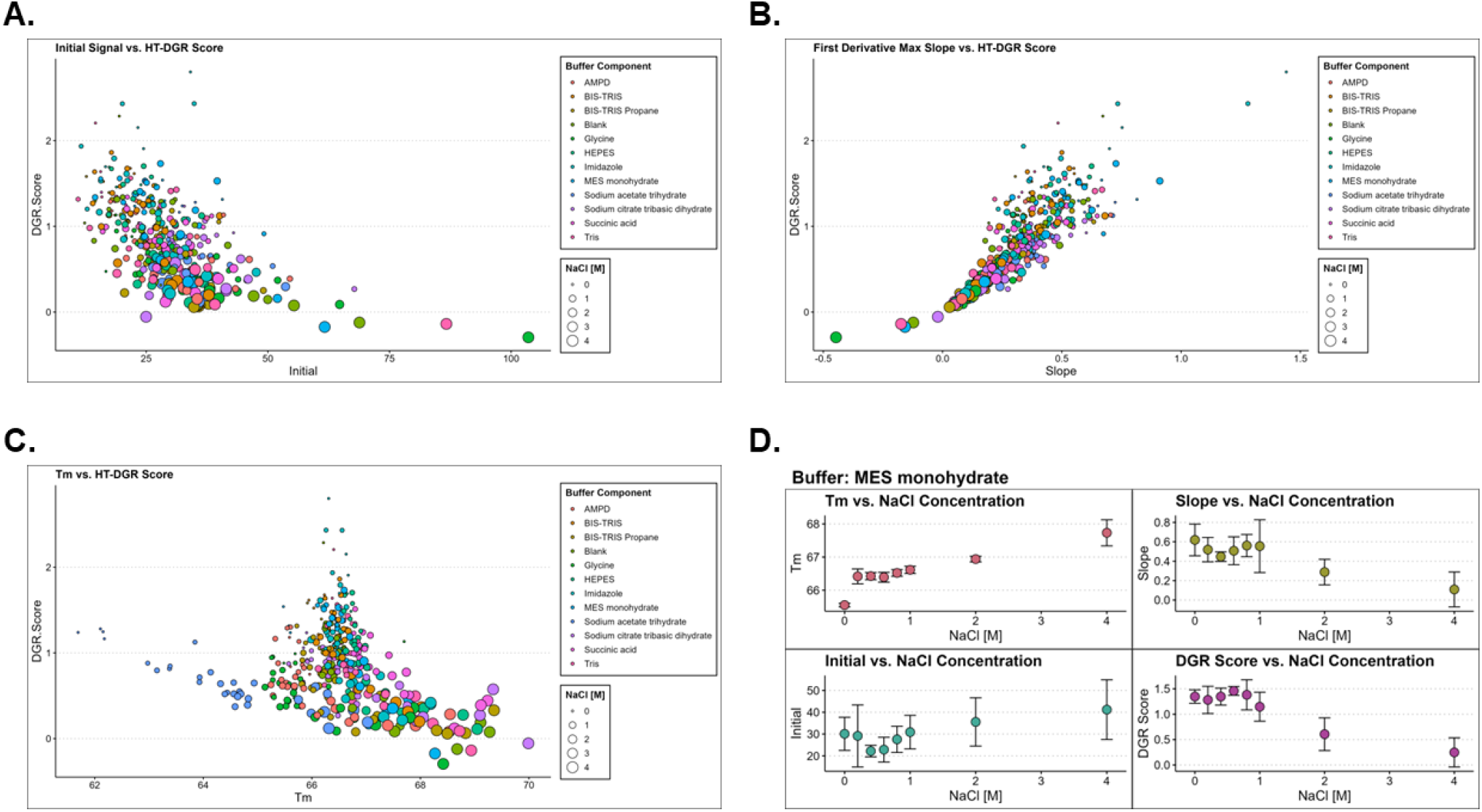
(A-C) Output of the DSF thermogram calculated parameters from the HT-DGR R script used to process thermal shift analysis data. Points are colored by the buffer component and sized according to the NaCl concentration. (D) Outputs for each of the parameters from the thermogram and DGR score are given for each buffer component against the NaCl concentration. Data presented is the mean of 4 replicates with error bars showing standard deviations.

## References

Amadeo, B. B., H. d. l. P. Andres, T. Roopa, T. Roopa, T. Roopa, Z. L. Jean and J. P. Kevin (2016). “DSF Guided Refolding As A Novel Method Of Protein Production.” Scientific Reports.

Backert, S., S. Bernegger, J. Skorko-Glonek and S. Wessler (2018). “Extracellular HtrA serine proteases: An emerging new strategy in bacterial pathogenesis.” Cell Microbiol 20(6): e12845.

Bai, X. C., X. J. Pan, X. J. Wang, Y. Y. Ye, L. F. Chang, D. Leng, J. Lei and S. F. Sui (2011). “Characterization of the structure and function of Escherichia coli DegQ as a representative of the DegQ-like proteases of bacterial HtrA family proteins.” Structure 19(9): 1328–1337.

Baynes, B. M., D. I. C. Wang and B. L. Trout (2005). “Role of Arginine in the Stabilization of Proteins against Aggregation.” Biochemistry 44(12): 4919–4925.

Cho, H., Y. Choi, K. Min, J. B. Son, H. Park, H. H. Lee and S. Kim (2020). “Over-activation of a nonessential bacterial protease DegP as an antibiotic strategy.” Commun Biol 3(1): 547.

Clausen, T., C. Southan and M. Ehrmann (2002). “The HtrA Family of Proteases.” Molecular Cell 10(3): 443–455.

Coleman, J. L., A. Toledo and J. L. Benach (2016). “Borrelia burgdorferi HtrA: evidence for twofold proteolysis of outer membrane protein p66.” Mol Microbiol 99(1): 135–150.

de Regt, A. K., S. Kim, J. Sohn, R. A. Grant, T. A. Baker and R. T. Sauer (2015). “A conserved activation cluster is required for allosteric communication in HtrA-family proteases.” Structure 23(3): 517–526.

Fox, B. G. and P. G. Blommel (2009). “Autoinduction of protein expression.” Current protocols in protein science Chapter 5: Unit-5.23.

François, B., B. François and M. Mirna (2004). “Recombinant protein folding and misfolding in Escherichia coli.” Nature Biotechnology.

Gherardini, F. C. (2013). “Borrelia burgdorferi HtrA may promote dissemination and irritation.” Mol Microbiol 90(2): 209–213.

Hansen, G. and R. Hilgenfeld (2013). “Architecture and regulation of HtrA-family proteins involved in protein quality control and stress response.” Cell Mol Life Sci 70(5): 761–775.

Jiang, J., X. Zhang, Y. Chen, Y. Wu, Z. H. Zhou, Z. Chang and S.-F. Sui (2008). “Activation of DegP chaperone-protease via formation of large cage-like oligomers upon binding to substrate proteins.” Proceedings of the National Academy of Sciences 105(33): 11939.

Kariu, T., K. Sharma, P. Singh, A. A. Smith, B. Backstedt, O. Buyuktanir and U. Pal (2015). “BB0323 and novel virulence determinant BB0238: Borrelia burgdorferi proteins that interact with and stabilize each other and are critical for infectivity.” J Infect Dis 211(3): 462–471.

Kariu, T., X. Yang, C. B. Marks, X. Zhang and U. Pal (2013). “Proteolysis of BB0323 results in two polypeptides that impact physiologic and infectious phenotypes in Borrelia burgdorferi.” Mol Microbiol 88(3): 510–522.

Kim, S., R. A. Grant and R. T. Sauer (2011). “Covalent linkage of distinct substrate degrons controls assembly and disassembly of DegP proteolytic cages.” Cell 145(1): 67–78.

Kolmar, H., P. R. Waller and R. T. Sauer (1996). “The DegP and DegQ periplasmic endoproteases of Escherichia coli: specificity for cleavage sites and substrate conformation.” Journal of Bacteriology 178(20): 5925–5929.

Krakowiak, J., M. Krajewska and J. Wawer (2019). “Monitoring of lysozyme thermal denaturation by volumetric measurements and nanoDSF technique in the presence of N-butylurea.” J Biol Phys 45(2): 161–172.

Krojer, T., J. Sawa, E. Schafer, H. R. Saibil, M. Ehrmann and T. Clausen (2008). “Structural basis for the regulated protease and chaperone function of DegP.” Nature 453(7197): 885–890.

Li, Z., W. Kessler, J. van den Heuvel and U. Rinas (2011). “Simple defined autoinduction medium for high-level recombinant protein production using T7-based Escherichia coli expression systems.” Appl Microbiol Biotechnol 91(4): 1203–1213.

Magnusson, A. O., A. Szekrenyi, H. J. Joosten, J. Finnigan, S. Charnock and W. D. Fessner (2019). “nanoDSF as screening tool for enzyme libraries and biotechnology development.” FEBS J 286(1): 184–204.

Pallen, M. J. and B. W. Wren (1997). “The HtrA family of serine proteases.” Mol Microbiol 26(2): 209–221.

Russell, T. M., M. J. Delorey and B. J. Johnson (2013). “Borrelia burgdorferi BbHtrA degrades host ECM proteins and stimulates release of inflammatory cytokines in vitro.” Mol Microbiol 90(2): 241–251.

Russell, T. M., X. Tang, J. M. Goldstein, D. Bagarozzi and B. J. Johnson (2016). “The salt-sensitive structure and zinc inhibition of Borrelia burgdorferi protease BbHtrA.” Mol Microbiol 99(3): 586–596.

Schubert, A., R. Wrase, R. Hilgenfeld and G. Hansen (2015). “Structures of DegQ from Legionella pneumophila Define Distinct ON and OFF States.” J Mol Biol 427(17): 2840–2851.

Thakur, M., S. Bista, S. D. Foor, S. Dutta, X. Yang, M. Ronzetti, V. S. Rana, C. Kitsou, S. B. Linden, A. S. Altieri, B. Baljinnyam, D. C. Nelson, A. Simeonov and U. Pal (2022). “Controlled Proteolysis of an Essential Virulence Determinant Dictates Infectivity of Lyme Disease Pathogens.” Infect Immun 90(5): e0005922.

Thompson, N. J., M. Merdanovic, M. Ehrmann, E. van Duijn and A. J. Heck (2014). “Substrate occupancy at the onset of oligomeric transitions of DegP.” Structure 22(2): 281–290.

Ullmann, A. J., T. M. Russell, M. C. Dolan, M. Williams, A. Hojgaard, Z. P. Weiner and B. J. Johnson (2015). “Evaluation of Borrelia burgdorferi BbHtrA Protease as a Vaccine Candidate for Lyme Borreliosis in Mice.” PLoS One 10(6): e0128868.

Wrase, R., H. Scott, R. Hilgenfeld and G. Hansen (2011). “The Legionella HtrA homologue DegQ is a self-compartmentizing protease that forms large 12-meric assemblies.” Proc Natl Acad Sci U S A 108(26): 10490–10495.

Ye, M., K. Sharma, M. Thakur, A. A. Smith, O. Buyuktanir, X. Xiang, X. Yang, K. Promnares, Y. Lou, X. F. Yang and U. Pal (2016). “HtrA, a Temperature-and Stationary Phase-Activated Protease Involved in Maturation of a Key Microbial Virulence Determinant, Facilitates Borrelia burgdorferi Infection in Mammalian Hosts.” Infect Immun 84(8): 2372–2381.

Yuanze, W., O. Niels van, M. A. Ameena, A. Alaa, L. A. Atsarina, D. Alexander and R. G. Matthew (2017). “A Systematic Protein Refolding Screen Method using the DGR Approach Reveals that Time and Secondary TSA are Essential Variables.” Scientific Reports.

Zarzecka, U., A. Grinzato, E. Kandiah, D. Cysewski, P. Berto, J. Skorko-Glonek, G. Zanotti and S. Backert (2020). “Functional analysis and cryo-electron microscopy of Campylobacter jejuni serine protease HtrA.” Gut Microbes 12(1): 1–16.

Zarzecka, U., A. Harrer, A. Zawilak-Pawlik, J. Skorko-Glonek and S. Backert (2019). “Chaperone activity of serine protease HtrA of Helicobacter pylori as a crucial survival factor under stress conditions.” Cell Commun Signal 17(1): 161.

Zarzecka, U., D. Matkowska, S. Backert and J. Skorko-Glonek (2021). “Importance of two PDZ domains for the proteolytic and chaperone activities of Helicobacter pylori serine protease HtrA.” Cell Microbiol 23(4): e13299.

Zarzecka, U., A. Modrak-Wojcik, M. Bayassi, M. Szewczyk, A. Gieldon, A. Lesner, T. Koper, A. Bzowska, M. Sanguinetti, S. Backert, B. Lipinska and J. Skorko-Glonek (2018). “Biochemical properties of the HtrA homolog from bacterium Stenotrophomonas maltophilia.” Int J Biol Macromol 109: 992–1005.

Zhang, K., Z. Qin, Y. Chang, J. Liu, M. G. Malkowski, S. Shipa, L. Li, W. Qiu, J. R. Zhang and C. Li (2019). “Analysis of a flagellar filament cap mutant reveals that HtrA serine protease degrades unfolded flagellin protein in the periplasm of Borrelia burgdorferi.” Mol Microbiol 111(6): 1652–1670.

Zhang, L., X. Wang, F. Fan, H. W. Wang, J. Wang, X. Li and S. F. Sui (2017). “Cryo-EM structure of Nma111p, a unique HtrA protease composed of two protease domains and four PDZ domains.” Cell Res 27(4): 582–585.

