## Supplementary figures and images for "Application of biophysical methods for improved protein production and characterization: a case study on an HtrA-family bacterial protease"

### Supplemental Figures

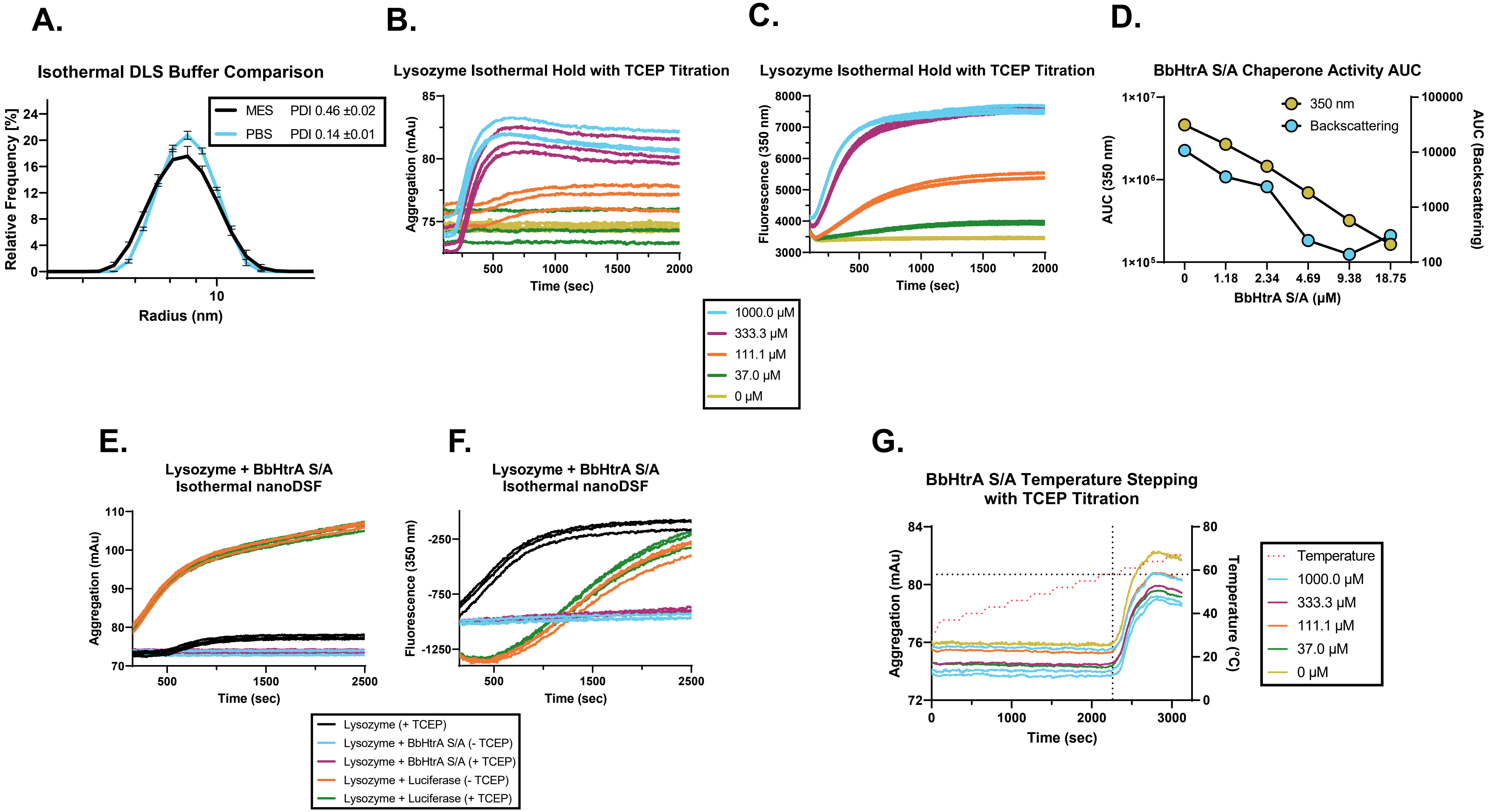

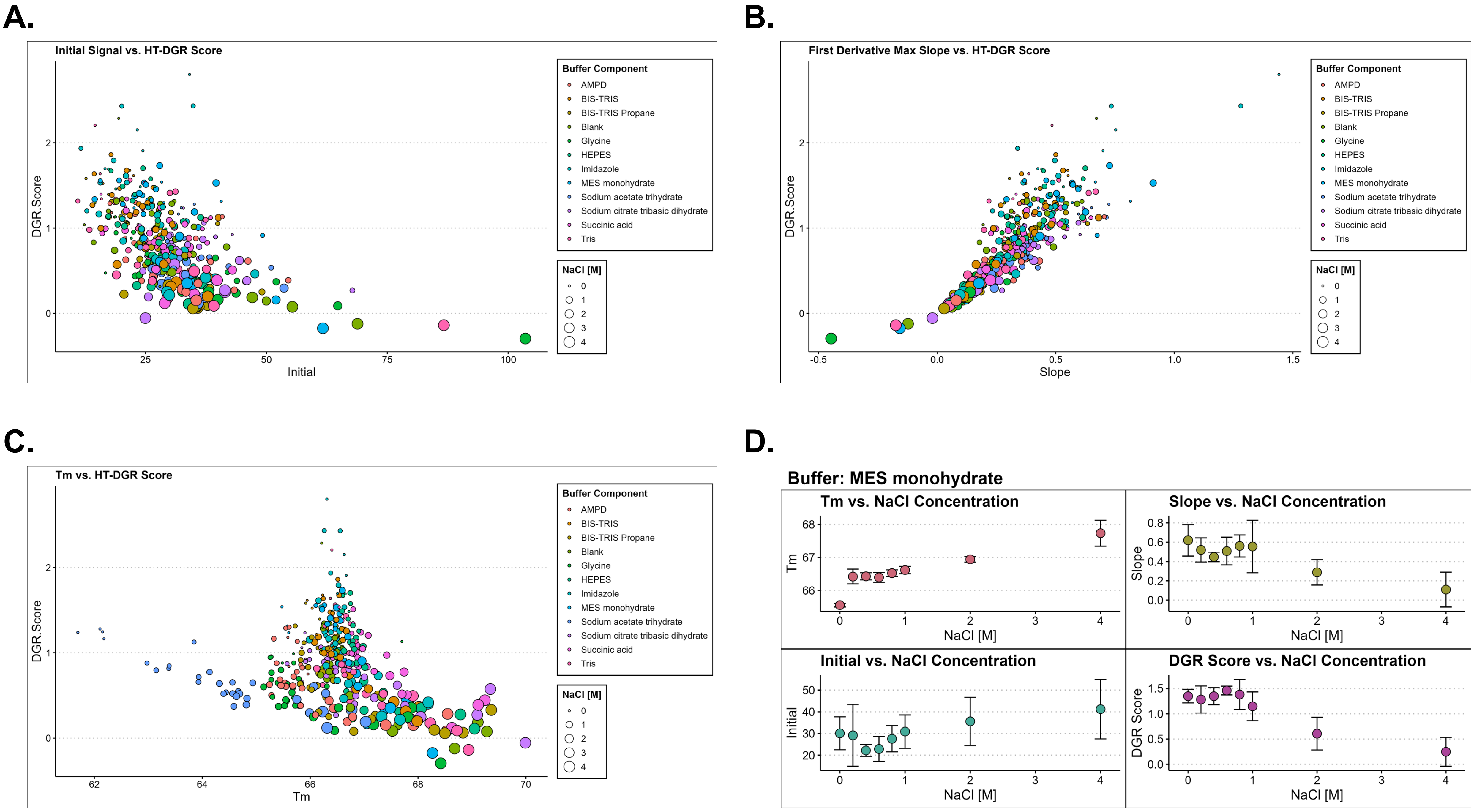
